# Computational Insight into Anti-Obesity Effects of Combined Some Phytobiotics to GLP1R (Glucagon-Like Peptide 1 Receptor) Protein in *Anas javanica*

**DOI:** 10.1101/2023.04.26.538390

**Authors:** CA Damayanti, MAY Harahap, S Wibowo, O Sjofjan, IH Djunaidi

## Abstract

Damayanti CA, Harahap MAY, Wibowo S, Sjofjan O, Djunaidi IH. 2023. Computational Insight into Anti-Obesity Effects of Indonesian Phytobiotics to GLP1R (Glucagon-Like Peptide 1 Receptor) Protein in *Anas javanica*.

Mojosari ducks (*Anas javanica*) is native Indonesia laying ducks was a egg producing type with quite high egg production, must be maintain body weight to propotional condition as laying duck. If the body weight surpasses normal, it can lead to obesity and reduce the eggs quality. One of the proteins closely related to obesity and hyperglycemia is GLP1R (Glucagon-Like Peptide 1 Receptor). The increase in GLP1R activity by one of the compounds that have been widely researched is loureirin B. Interaction between loureirin B and GLP1R increases insulin production in the body so that hyperglycemia and body weight can be controlled properly. Exploration of phytobiotic compounds from Indonesia is needed to find the substitution of loureirin B as an anti-obesity agent. According to the findings of in silico study (protein modeling and molecular docking), cynaroside (−9.2 kcal/mol), 14-Deoxy-11,12-didehydroandrographolide (−9.1 kcal/mol), rutin (−8.8 kcal/mol), andrographidine E (−8.6 kcal/mol), and cianidanol (−7.8 kcal/mol) had stronger binding affinity than loureirin B (−7.4 kcal/mol). Andrographidine E, derived from the plant *Andrographis paniculata*, is the best candidate for GLP1R agonist. The binding affinity that Andrographidine E has is lower than control compounds, so it is easier for bonds to occur between proteins and such compounds. In addition, the interacting amino acids do not have unfavourable bonds that make it more stable than other candidates. Results from clinical studies show that the use of *A. paniculata* can reduce glucose levels.

## INTRODUCTION

It is known that the parameters of the success of a livestock business are inseparable from the role of feed. In order to maximize the nutritional value of feed consumed by poultry, the use of feed additives as a feed ingredient that is added with a small amount to the feed is vital. Two types of feed additives can be added to feed: natural and synthetic. In modern poultry farms, synthetic feed additives such as antibiotics (Antibiotics Growth Promoters / AGP) are generally used to increase feed efficiency, spur growth, and increase productivity. However, the use of antibiotics as feed additives is feared to cause antibiotic residues that can cause bacterial resistance, so it has a harmful impact on livestock and the health of the humans who consume them. The approach of giving natural feed additives derived from spice plants began to be widely used as an alternative to antibiotics. As an agricultural country, Indonesia is rich in biodiversity, with fertile land and land that supports the agricultural sector as the second most significant support for the Indonesian economy. One of the reliable products from the agricultural sector is spices containing phytochemical compounds produced by plants as part of metabolic processes. Exploration of the potential of spices from Indonesia such as *Curcuma longa, Zingiber officinale, Kaempferia galanga, Andrograpis paniculate, Tinospora crispa, Phyllanthus niruri*, and *Piper betle Linn* are currently reported to have an essential role as a natural feed additive in maintaining health, improving production appearance, and increasing laying poultry productivity.

Mojosari ducks (*Anas Javanica*) is native Indonesia laying ducks was a egg producing type with quite high egg production, must be maintain body weight to propotional condition as laying duck. If the body weight exceeds normal, it can trigger obesity, where fat deposits on the abdomen can reduce the elasticity of the oviduct because fat piles restrain it. So that when contractions occur, the oviduct will be difficult to return to its original position and even allow some of the eggs to be outside. This condition triggers cases of prolapse, which has an impact on reducing the productivity of ducks in producing eggs (Ray et al. 2013). The presence of hyperglycemia characterizes a common metabolic disease in laying birds as a result of a decrease in insulin secretion. The mechanism of insulin secretion in the body is inseparable from the role of the hormone incretin as a stimulant of insulin secretion, one of which is the GLP1 protein (Glucagon-Like Peptide 1) (Sumaryada et al. 2021) by activating the GLP1R (Glucagon-Like Peptide 1 Receptor) agonist in cells β the pancreas (Meloni et al. 2012). So to suppress glycemic levels in the blood, it is necessary to increase the activity of GLP1R by developing the GLP1 protein agonist to maintain the amount of GLP1 in the body. One of the agonists that have been proven to perform an interaction with GLP1R is Loureirin B. Loureirin B is a natural compound derived from *Sanguis draconis* and has various pharmacological effects, one of which is hypoglycemic. The interaction in Loureirin B is similar to the interaction of the GLP1 protein, which interacts with GLP1R to increase insulin secretion (Ding et al. 2021).

Through the in silico approach, these spices’ active compounds can be studied as a substitute for Loureirin B in increasing insulin secretion. Research using active compounds from various spices in Indonesia and its relation to the GLP1R protein in Mojosari ducks has never been carried out in silico.

## MATERIALS AND METHODS

### Data collection

To obtain data in in-silico studies, control ligands and phytobiotic data were obtained from the PubChem database. The control ligands used were loureirin B (189670), and the phytobiotics used were gingerol (442793), tyramine (5610), rutin (5280805), 14-deoxy-11,12-didehydroandrographiside (44575271), andrographidine-e (13963769) (969516). Meanwhile, information for protein modeling was collected from Uniprot for proteins with the code B4ZY91 (Glucagon-Like Peptide 1 Receptor) is a protein from *Gallus gallus*, which contains a maximum of 459 amino acids (Wibowo et al. 2022). The structure of the proteins and ligands can be seen in Table 1.

**Table 1.**
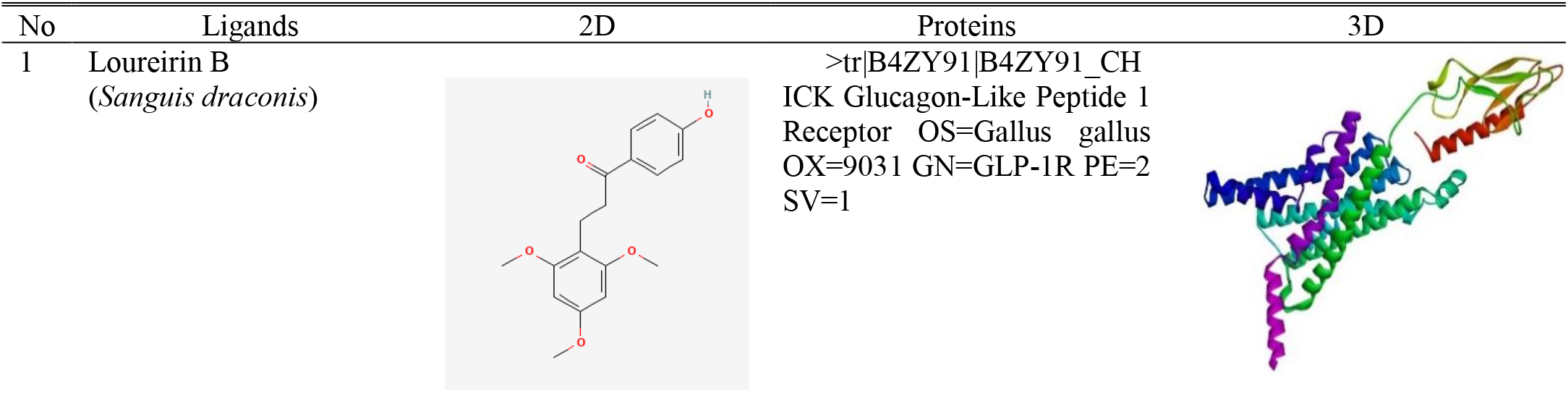
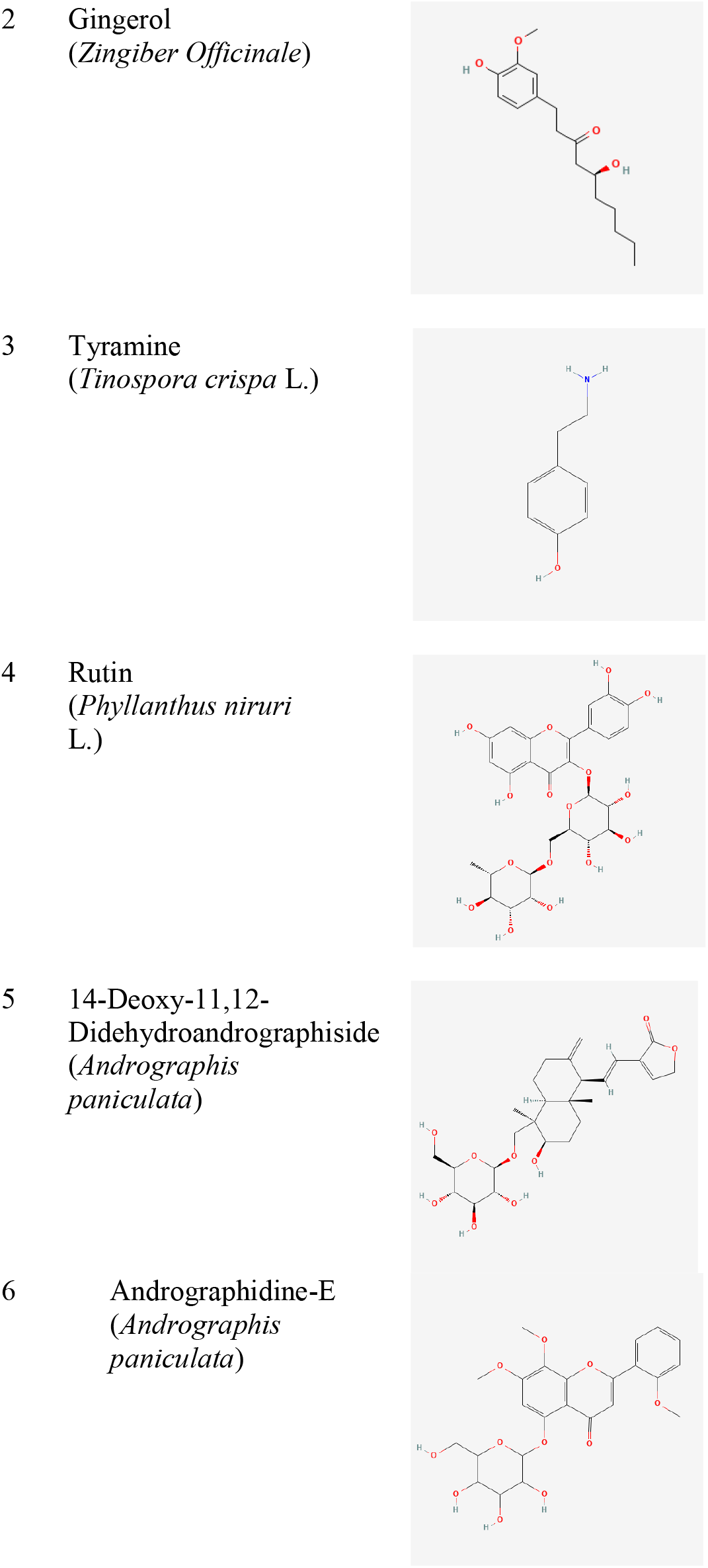
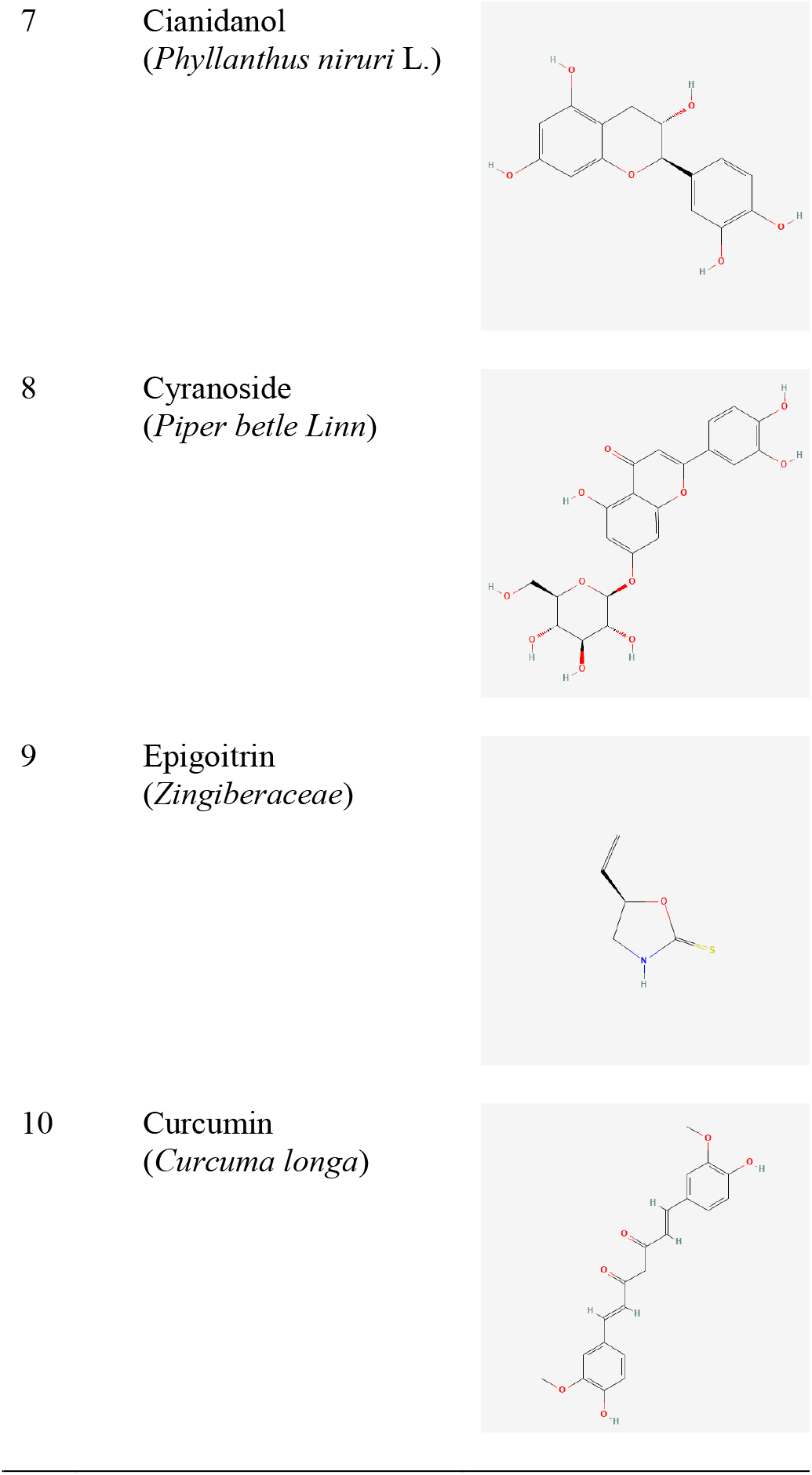
Structure of the proteins and ligands

### Protein modelling and assessment

SWISS-MODEL protein modeling with input of protein amino acid sequences. Model selection based on results from the protein template, GMQE, QSGE, and identification tests. After the protein has been correctly modeled, RAMPAGE is used in the evaluation process to determine the protein’s quality based on the Ramachandran Plot (Komari et al. 2020).

### Molecular Docking

The Discovery Studio 2016 Client was used for protein preparation. Proteins are separated from water molecules and their ligands. The ligands are then prepared in Open Babel for conversion (.sdf) to (.pdbqt). Furthermore, furthermore, docking uses Autodock Vina which is integrated with CB-Dock 2 (https://cadd.labshare.cn/cb-dock2/php/index.php) (Liu et al. 2022). The docking results with the highest rank and passed the validation (RMSD ≤ 2 Å). Data docking analysis and visualization with Discovery Studio 2016 Client (Wibowo et al. 2020) (Wibowo et al. 2019) (Wibowo et al. 2021).

## RESULTS AND DISCUSSION

### Protein Modelling and Assesement

The three-dimensional (3D) structure largely determines the nature and function of proteins biochemically. In contrast, determining the 3D structure of proteins can be done quickly and cheaply through the in silico method. The 3D structure modeling method of proteins is divided into three methods, namely homology modeling, fold recognition, and ab initio. Homology modeling is the best 3D structure modeling of proteins whose modeling is carried out by aligning the amino acid sequence of the target protein with other proteins that have been instrumentally known as 3D structures. Proteins already known to be 3D structures are referred to as templates (Komari et al, 2020). Protein structure modeling uses a SWISS-MODEL server because it has a high degree of efficiency, so the GLP1R protein can be used for further analysis. SWISS-MODEL is used by following several stages, including determining protein target sequences, identifying protein templates, making models, and evaluating models (Komari et al, 2020). The protein template used in this study refers to 7duq.1.B, which is the Glucagon-Like Peptide 1 Receptor. Template selection is based on coverage value, global model quality estimation of 0.73, and identity, which reaches 78.39. The following is a visualization of the protein modeling presented in Figure 1.

**Figure 1.**
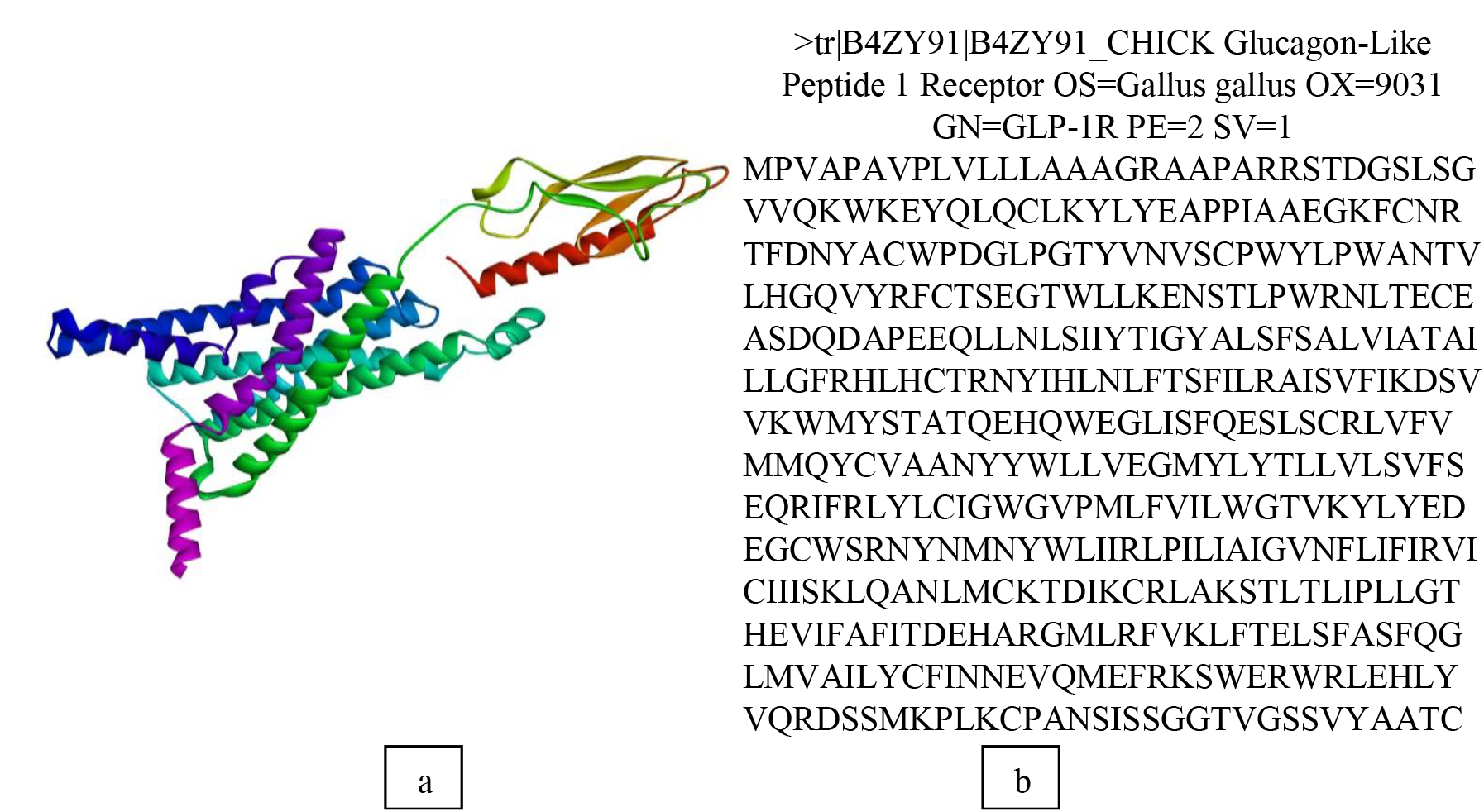
Visualization of the protein modelling Description a. Protein Modelling GLP-1R b. Amino Acid Sequence

Protein modelling data was obtained from Uniprot with the code B4ZY91 (Glucagon-Like Peptide 1 Receptor) with a total of 459 aa amino acids. The protein used for modeling is a protein from poultry (Gallus gallus). Glucagon-Like Peptide 1 is a type of incretin hormone produced by L cells in the ileum, plays a role in the mechanism of action of insulin as a stimulant of insulin secretion by activating the GLP1R agonist (Glucagon-Like Peptide 1 Receptor) in cells β the pancreas even though there are also cells α the pancreas. GLP-1 hormone will work when glucose concentration is above basal concentration (Muller et al, 2019). After protein modeling, it was continued with the evaluation of the results of homology modelling using the Procheck web server which is a program for validation of protein modelling by analyzing the Ramachandran plot. The Ramachandran plot is a two-dimensional plot that describes amino acid residues in enzyme structures that have been determined through experiments into internal coordinates, where the angle Φ (phi) is the x-axis while the ψ (psi) is the y-axis divided into four quadrudes. The quality assessment is based on the percentage of amino acid residues present in the most favoured regions and disallowed regions of the Ramachandran plot (Junaidin et al, 2019). Ramachandran’s plot for the GLP-1R (Glucagon-Like Peptide 1 Receptor) protein model presented in Figure 2.

**Figure 2.**
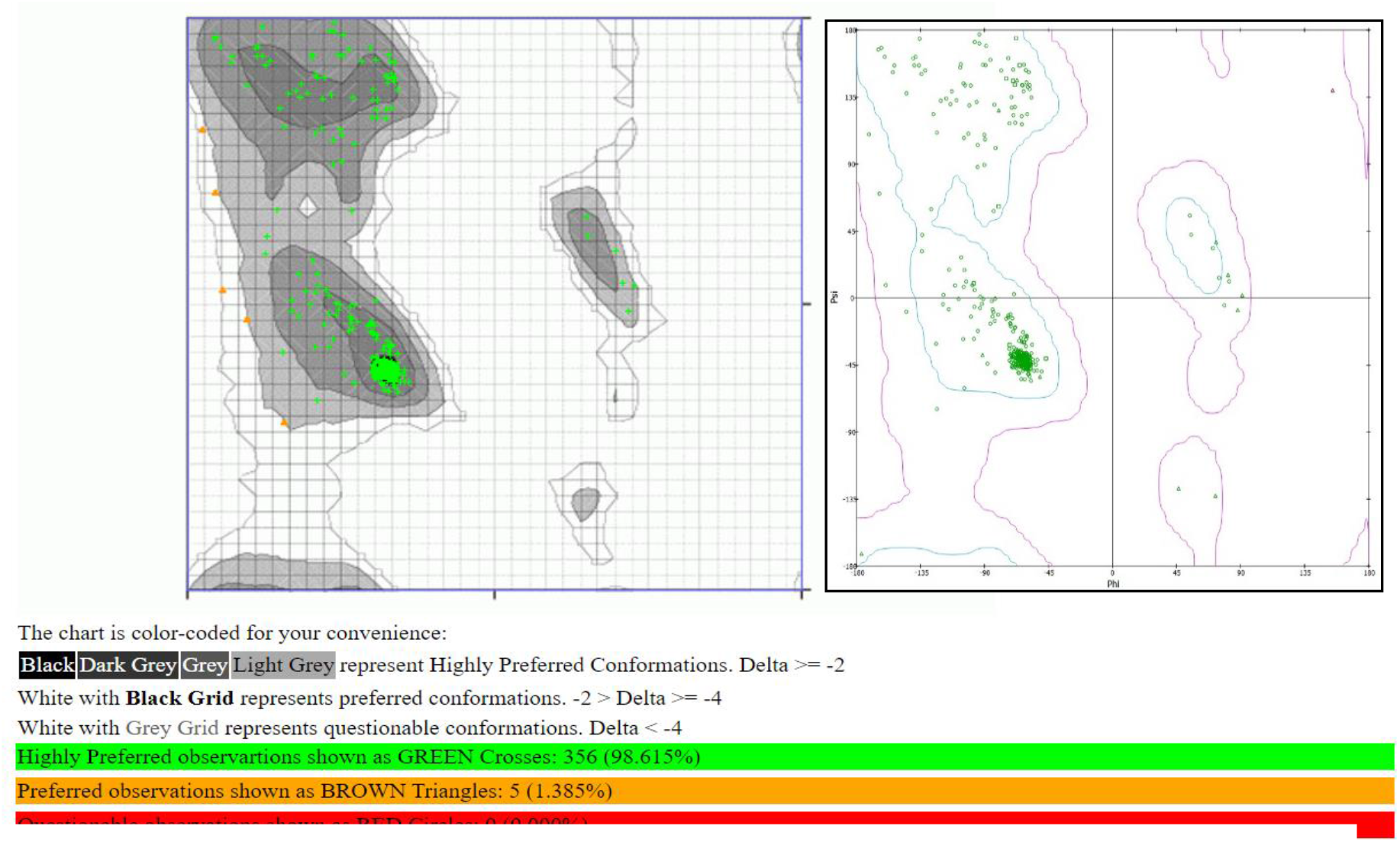
(Glucagon-Like Peptide 1 Receptor) protein model

### Molecular Docking

*Curcuma longa, Zingiber Officinale, Zingiberaceae, Andrograpis paniculata, Tinospora crispa* L., *Phyllanthus niruri* L. and *Piper betle Linn* which are components of phytobiotics have been known to have an important role as a natural feed additive in maintaining health, improving production appearance, and increasing poultry productivity. Through the in silico method approach, the active compounds in the spice were found nine active compounds that can be studied as GLP1R agonists in increasing insulin secretion. The active compounds are curcumin (*Curcuma longa*), ethyl-p-methoxycinnamte/epigoitrin (*Kaempferia galanga*), gingerol (*Zingiber Officinale*), cynaroside (*Piper betle Linn*), rutin and cianidnol (*Phyllanthus niruri*), andrographidine E and 14-Deoxy-11,12-didehydroandrographolide (*Andrograpis paniculata*), and tyramine (*Tinospora crispa*). The following are the results of molecular docking on active compounds in phytobiotics that have criteria as GLP1R agonists.

The lourerin B and GLP-1R complexes have several types of bonds, namely conventional hydrogen bonds with amino acids, namely PHE(B):363, GLU(B):383, and TYR(B):237. Then there is also a carbon hydrogen bond with amino acids, namely ARG (B):306 and TYR(B):144. Next is the pi-sulfur bond with the amino acid MET(B):229, as well as the pi-sigma bond in VAL(B):233 (Figure 3). The next step after obtaining the type of bond and protein in the control material, a candidate inhibitor of Loureirin B was sought in the active compound in the phytobiotics. The following are the results of molecular docking on active compounds in phytobiotics that have criteria as GLP-1R agonists.

**Figure 3.**
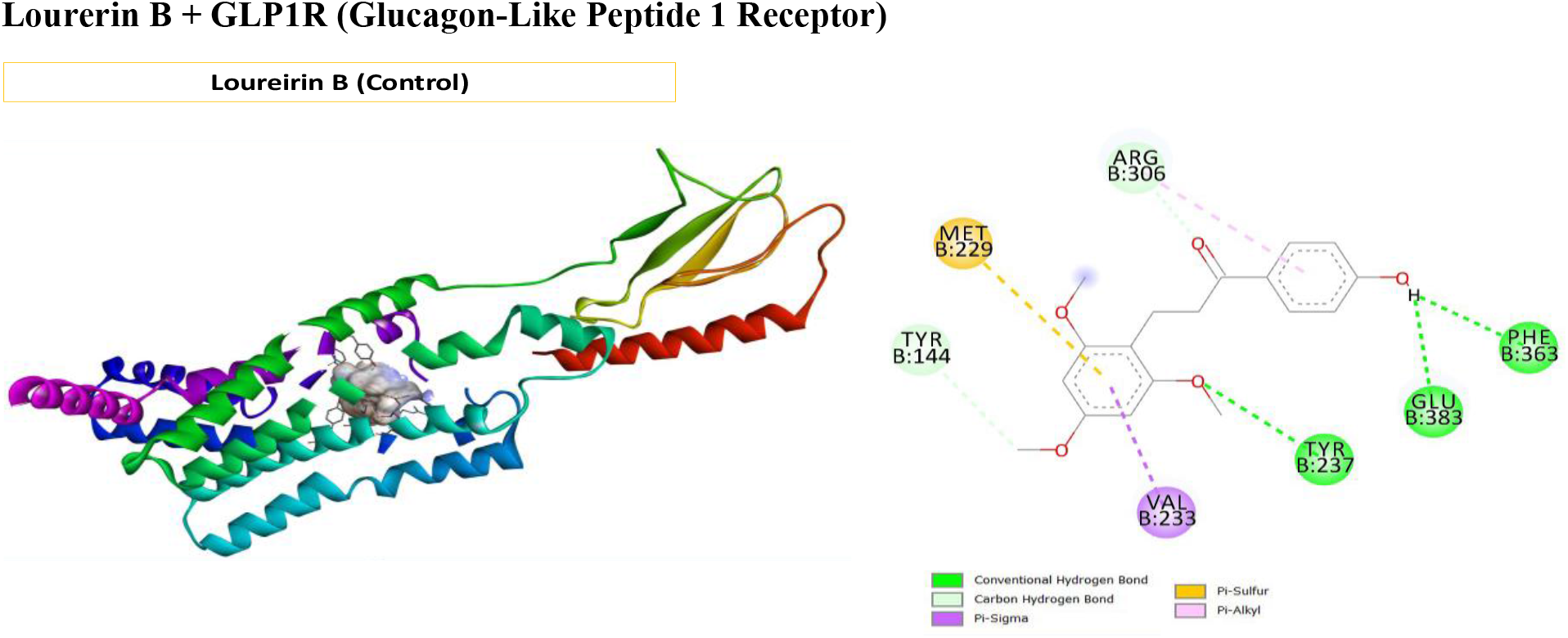
Visualization docking Lourerin B + Glucagon-Like Peptide 1 Receptor

Figure 4a conventional hydrogen bonds are found in the amino acids GLU(B):383, ILE(B):366, ASP(B):368, TYR(B):237, and LYS(B):193. The next bonds are pi-alkyl in the amino acids LEU(B):384, and alkyls in LEU(B):380, LEU(B):140, LEU(B):147, and VAL(B):233. And there is one bond that is unfavorable acceptors in GLU(B):360. Location of the amino acid rutin in the results of docking routine compounds consists of four large groups, namely TYR (B):237, TYR(B):144, ARG(B):376, ASN(B):296, LYS(B):193, LYS(B):379, SER(B):294 (conventional hydrogen bond), GLU(B):383 (pi-anion bond), LEU(B):384 (pi sigma), and ASP(B):368 (unfavorable acceptor-acceptor bond) (Figure 4b). Conventional hydrogen bonds in the Andrographidine E can be found in TYR(B):237, GLU(B):360, ILE(B):366, and LYS(B):379. Then the carbon hydrogen bond is GLN(B):230 and pi-alkyl in LEU(B):280. Next up are pi-cation bonding in ASP(B):368 and ARG(B):376 as well as pi-pi t-shaped in TRP(B):302 and unfavorable positive-positive in LYS(A):727 (Figure 4c). In conventional hydrogen bond bonds found in the cianidanol and GLP1R complexes in the amino acids ALA(B):364, ASP(B):368, LYS(B):379, GLU(B):360 and GLN(B):230. Meanwhile, carbon hydrogen bonds in THR(B):367 and pi-alkyl bonds in LEU(B):310 and ARG(B):306. The pi-anion bond is on the amino acid GLU(B):383, then the pi-sulfur bond in MET(B):229 and the pi-sigma bond in VAL(B):233 (Figure 4d). There are three large bond groups from docking results in cynaroside, namely conventional hydrogen bonds, sigma bonds, and donor-donor unfavorable bonds. Conventional hydrogen bond bonds are found in the amino acids ARG(B):186, LYS(B):379, PHE(B):363, ILE(B):366, ASN(B):296. Sigma bonds were found in the amino acids MET(B):229 then bond GLN(B): 230 (Figure 4e).

**Figure 4.**
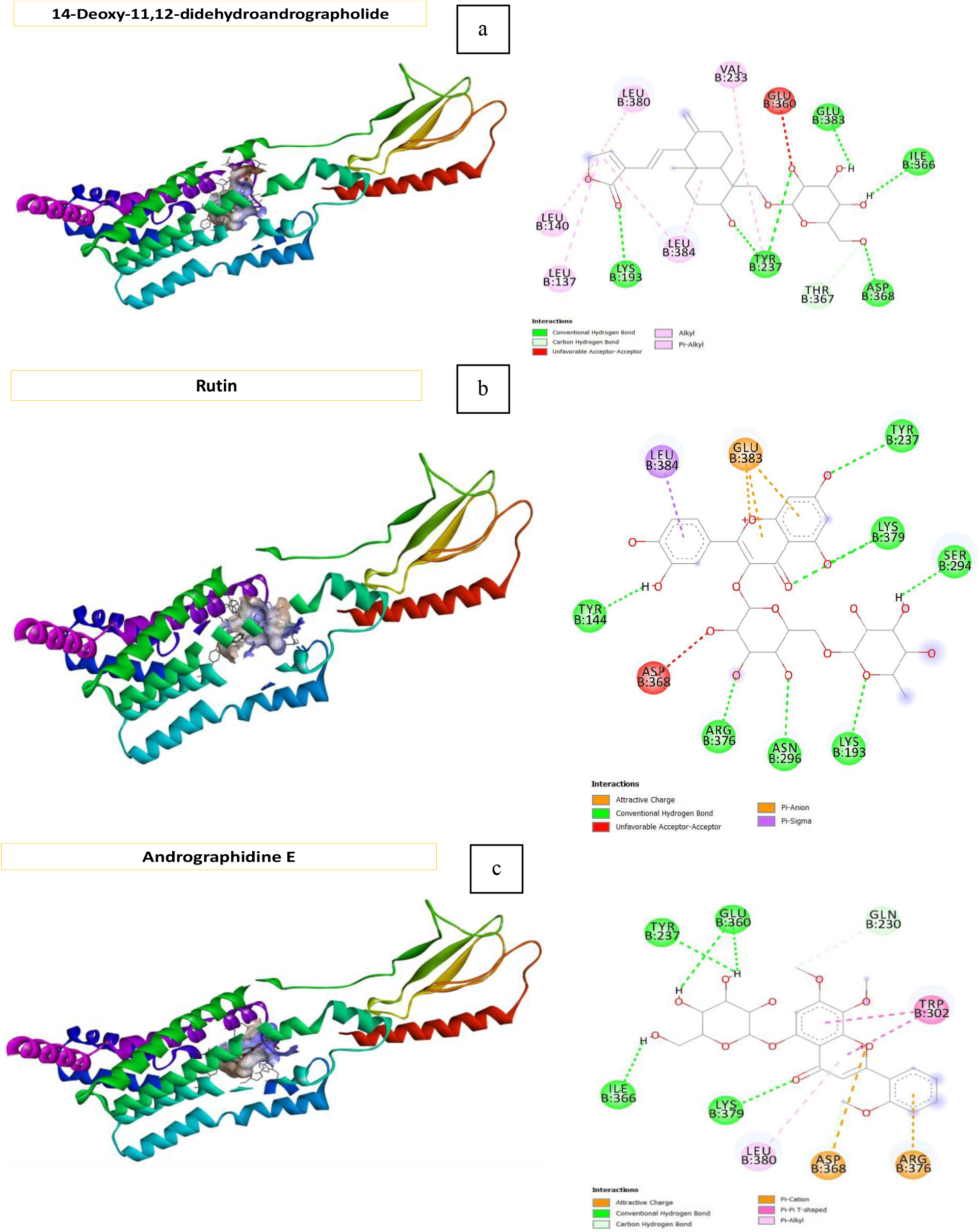
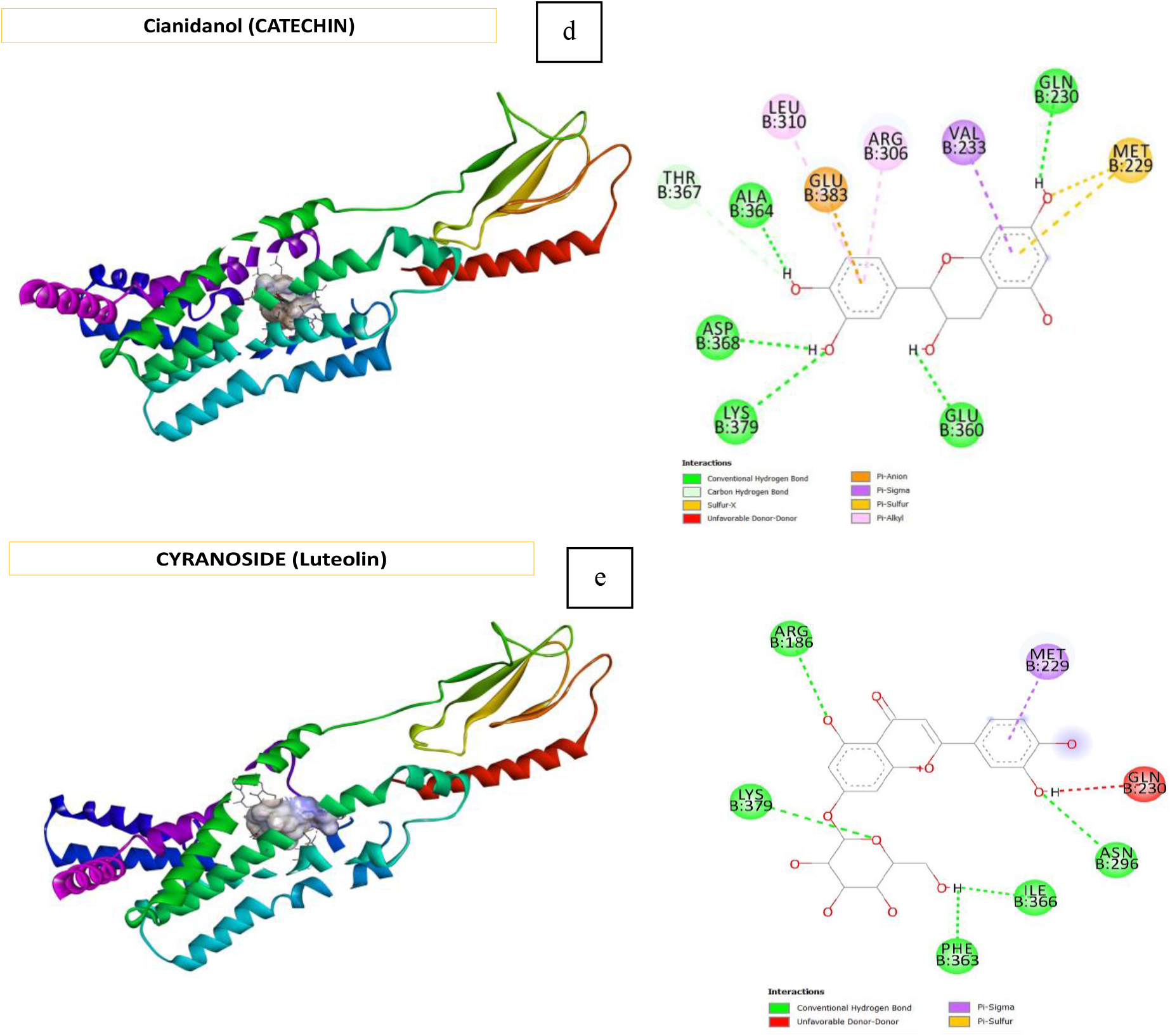
Visualization docking Description a. GLP-1R+*Rutin*, b. GLP-1R+*14-Deoxy*, c. GLP-1R *Andro E*, d.GLP-1R *Cianidanol*, e. GLP1+*Cynaroside*

Docking results on gingerol, tyramine, epigoitrin, and curcumin compounds that bind to GLP1R show that these compounds have a low probability as GLP1R agonists and Loureirin B inhibitors. The docking visualization can be seen in Figure 5 below.

**Figure 5.**
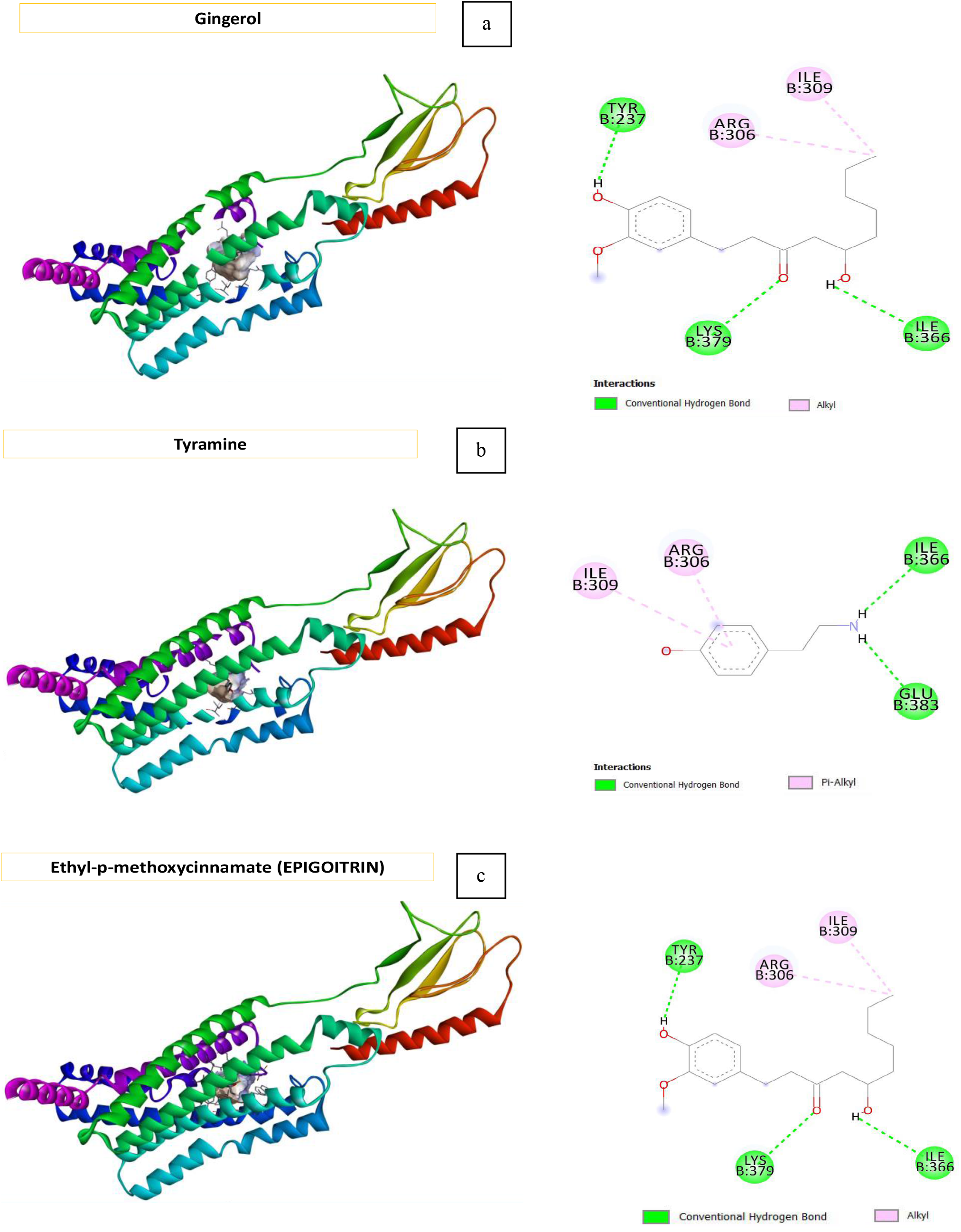
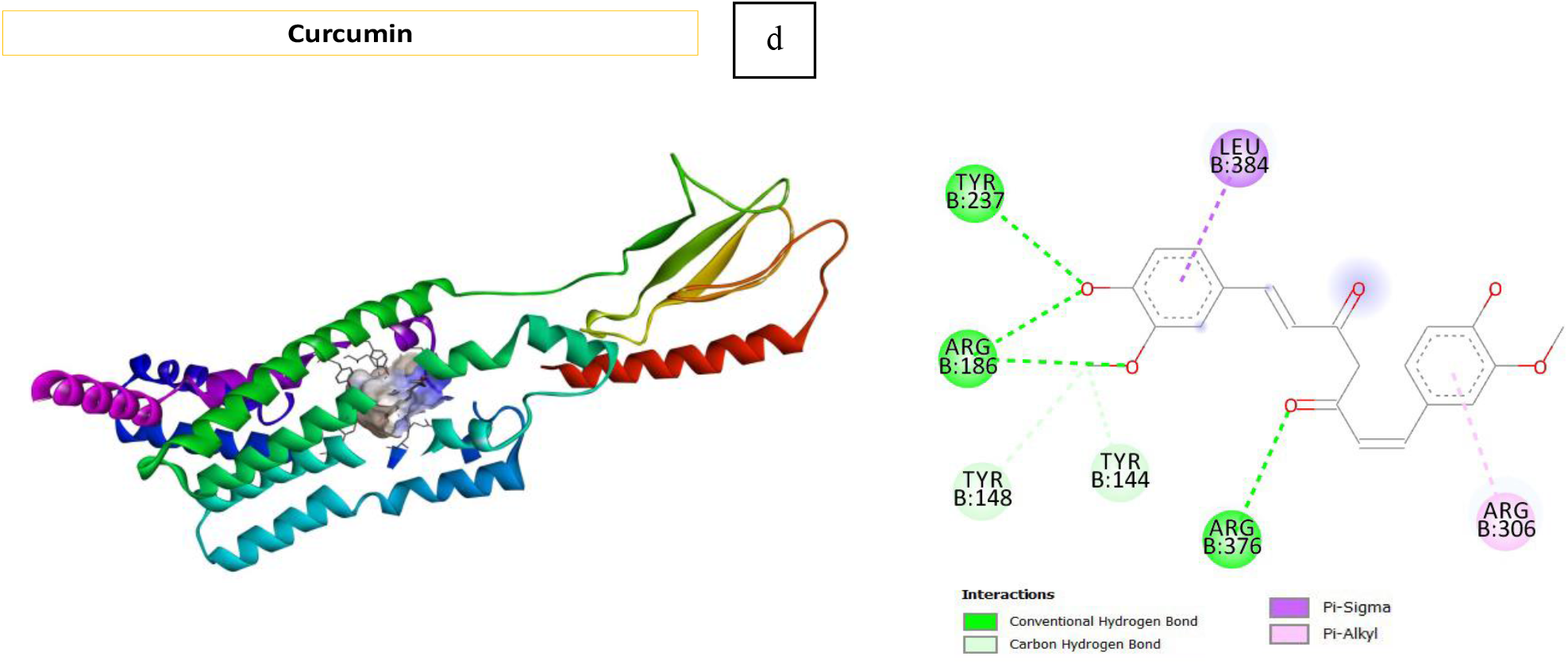
Visualisasi docking Description: a. GLP-1R+*Gingerol*, b. GLP-1R+ *Tyramine*, c. GLP-1R+ *Epigoitrin*, d. GLP-1R+ *Curcumin*

The docking results of Gingerol compounds have two large bond locations consisting of TYR(B):237, LYS(B):379, ILE(B):366 (conventional hydrogen bond bond) and ARG(B):306, ILE(B):309 (alkyl bond) (Figure 5a). In Figure 5b conventional hydrogen bonds are found in the amino acids ILE(B):366 GLU(B):383 and pi-alkyl bonds are found in the amino acids ARG(B):306, ILE(B):309. Conventional hydrogen bond bonds on the Epigoitrin and GLP1R complexes are found in the amino acids LYS(B):379, ASP(B):368, ALA(B):364. Then for alkyl bonds, it is found in the amino acids ILE(B):309 and ARG(B):306. Next for the Van der Waals bond on the amino acid ILE(B):366 ILE(B):313 THR(B):367 PHE(B):365 PHE(B):363 GLU(B):383 (Figure 5c). The location of the conventional hydrogen bond on curcumin and GLP1R is located in the amino acids ARG(B):186, ARG(B):376, TYR(B):237. Carbon hydrogen bonds are located in the amino acids TYR(B):144, TYR(B):148. The pi-alkyl bond is located in the amino acid ARG(B):306. Then the pi-sigma bond lies in the amino acid LEU(B): 384 (Figure 5d).

In this study, the use of homology modeling is a tool for predicting protein structure in three dimensions. The SWISS-MODEL used has a high degree of efficiency, so the GLP1R protein can be used for further analysis (Pitman et al, 2006) (Vyas et al, 2012) (Schwede et al, 2003). The protein template used in this study refers to 7duq.1.B, a Glucagon-Like Peptide 1 Receptor. Template selection is based on coverage value, global model quality estimation of 0.73, and identity, which reaches 78.39. The docking process of the proteins selected from the modeling results was then identified with control compounds, namely loureirin B and nine other phytobiotic compounds. The result of the interaction between loureirin B and GLP1R has a bond energy of −7.4 kcal/mol. It was previously known that loureirin B could increase insulin production by interacting with the GLP1R protein (Ding et al, 2021). When compared to the docking results of various other active compounds, lourerin B is still at fairly weak bond energy when compared to cynaroside (−9.2 kcal/mol), 14-Deoxy-11,12-didehydroandrographolide (−9.1 kcal/mol), rutin (−8.8 kcal/mol), andrographidine E (−8.6 kcal/mol), and cianidanol (−7.8 kcal/mol).

Meanwhile, other compounds such as curcumin, gingerol, tyramine, and epigoitrin have lower binding energy than controls (Figure 6). The research of Van et al, (2021) shows that cyranoside is proven effective in reducing hyperglycemia (Ding et al, 2021). Compared to cynaroside control, there are five hydrogen bonds, while only three controls have unfavorable donor-donor bonds. Likewise, the compound 14-Deoxy-11,12-didehydroandrographolide and rutin have five and seven hydrogen bonds, but the interaction of the two compounds with GLP1R has unfavorable bonds. Unfavorable bonds will generally have instability when testing molecular dynamics (Wibowo et al, 2021). The Andrographidine E compound is the best candidate as agonist for GLP1R. This is because the binding affinity is lower than control compounds, so it is easier to bond between proteins and these compounds. In addition, the interacting amino acids do not have unfavorable bonds, which makes them much more stable than other candidates. Clinical studies show that using *A. paniculate* can reduce glucose levels and lower body mass index in people with diabetes mellitus (Novianto et al, 2018). *Andrograpis paniculate* is an herbal plant in Indonesia that grows wildly, producing andrographolide compounds that produce a bitter taste, so it is known as the King of bitters. This in silico study found that andrographidine E can bind to the GLP1R protein and has the same bond site as loureirin B control, namely in TYR(B):237, and by having more hydrogen bonds than control, makes it more stable.

**Figure 6.**
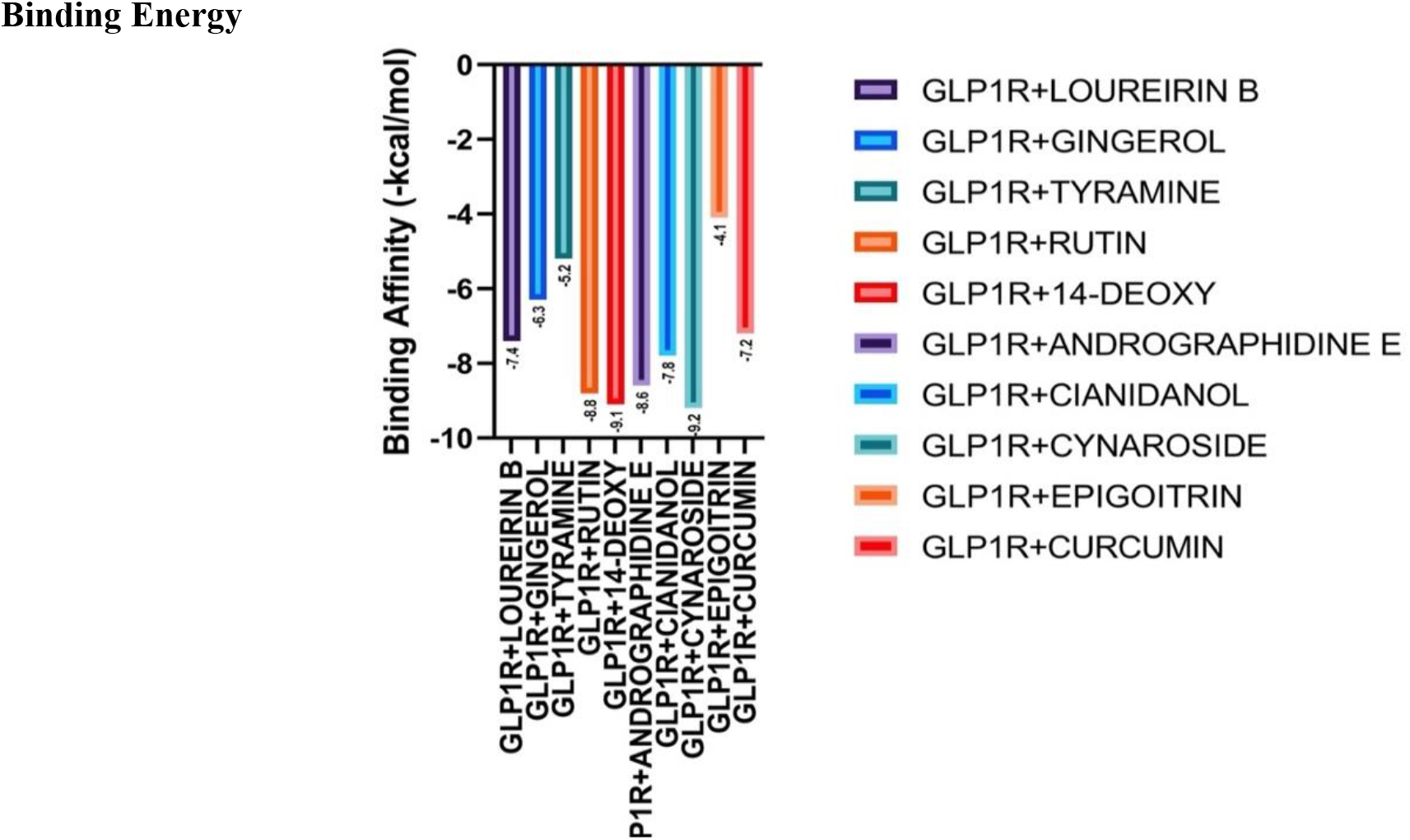
Binding Energy. Binding affinity from all of active compounds with GLP-1R showed in Figure 6. Compound that have best binding affinity is cymaroside. The weakest binding affinity showed in epigoitrin complex with GLP1R (−4.1 kcal/mol).

Further research, both in vitro and in vivo, is being conducted in our lab. Some other studies showed that analogs andrographolide and 14-Deoxy-11,12-didehydroandrographolide, including andrographidine E, have the antidiabetic ability through protection pathways on beta cells (Lee et al, 2021). This study shows new information that andrographidine E can be an agonist on the GLP1R protein. Andrographolide analogs also increased glucose uptake in 3T3-L1 cells by targeting TNF-a through NF-kB pathway activation. Activation of the pathway involves PI3-K (phosphatidylinositol 3-kinase), thus improving resistance to insulin (Zhang et al, 2009). 14-Deoxy-11,12-didehydroandrographolide (deAND) is included in the bioactive component of *A. paniculata*, which has antidiabetic activity. AMP-activated protein kinase (AMPK) regulates glucose transport and improves insulin resistance (Cheng et al, 2021).

Furthermore, rutin from Phyllanthus niruri L. is also known to carry out glycemic control levels through increased GLP1 levels in the body (Lee et al, 2021). Through a molecular approach, curcumin demonstrates its role as an anti-diabetic and antioxidant involving nuclear factor-kappa B (NF-kB), PPARγ, and free fatty acids. Curcumin is one of the active compounds from Curcuma longa that will inhibit the NF-kB pathway by suppressing the inflammatory rate, thereby reducing the damage to pancreatic beta cells that fail to secrete insulin in response to an increase in blood glucose levels in the body so that the body experiences hyperglycemia. Targets on curcumin include gene expression, enzymes, inflammatory cytokines, receptors, protein kinase, growth factors, and transcription factors (Malik et al, 2021). Gingerol (*Zingiber officinale*) is also reported to be able to correct hyperglycemia by restoring disturbed endocrine signals through increased GLP1 plasma, thereby increasing insulin secretion (Samad et al, 2017).

## CONCLUSION

From this study it can be concluded that cynaroside, 14-Deoxy-11,12-didehydroandrographolide, rutin, andrographidine E, and cianidanol have better binding affinity than loureirin B. Best candidate as a GLP-1R agonist agent is andrographidine E from the plant *A. paniculata*. Binding affinity that Andrographidine E has lower than control compounds so that it is easier for bonds to occur between proteins and such compounds. In addition the interacting amino acids do not have unfavourable bonds that making it more stable than other candidates. Results from clinical studies show the use of *A. paniculata* can reduce glucose levels.

## ACKNOWLEDGEMENT

The authors would like to thank Department of Animal Nutrition, Faculty of Animal Science, Brawijaya University for kind support during this research.

